# Cortical dynamics of saccade-target selection during free-viewing of natural scenes

**DOI:** 10.1101/075929

**Authors:** Linda Henriksson, Kaisu Ölander, Riitta Hari

## Abstract

Natural visual behaviour entails explorative eye movements, saccades, that bring different parts of a visual scene into the central vision. The neural processes guiding the selection of saccade targets are still largely unknown. Therefore, in this study, we tracked with magnetoencephalography (MEG) cortical dynamics of viewers who were freely exploring novel natural scenes. Overall, the viewers were largely consistent in their gaze behaviour, especially if the scene contained any persons. We took a fresh approach to relate the eye-gaze data to the MEG signals by characterizing dynamic cortical representations by means of representational distance matrices. Specifically, we compared the representational distances between the stimuli in the evoked MEG responses with predictions based (1) on the low-level visual similarity of the stimuli (as visually more similar stimuli evoke more similar responses in early visual areas) and (2) on the eye-gaze data. At 50–75 ms after the scene onset, the similarity of the occipital MEG patterns correlated with the low-level visual similarity of the scenes, and already at 75–100 ms the visual features attracting the first saccades predicted the similarity of the right parieto-occipital MEG responses. Thereafter, at 100–125 ms, the landing positions of the upcoming saccades explained MEG responses. These results indicate that MEG signals contain signatures of the rapid processing of natural visual scenes as well as of the initiation of the first saccades, with the processing of the saccade target preceding the processing of the landing position of the upcoming saccade.

**SIGNIFICANCE STATEMENT:** Humans naturally make eye movements to bring different parts of a visual scene to the fovea where our visual acuity is the best. Tracking of eye gaze can reveal how we make inferences about the content of a scene by looking at different objects, or which visual cues automatically attract our attention and gaze. The brain dynamics governing natural gaze behaviour is still largely unknown. Here we suggest a novel approach to relate eye-tracking results with brain activity, as measured with magnetoencephalography (MEG), and demonstrate signatures of natural gaze behaviour in the MEG data already before the eye movements occur.

## INTRODUCTION

Natural visual behaviour involves eye movements that bring different objects to the central retina where the visual acuity is the best. During viewing of real-world scenes, one fixation lasts a few hundred milliseconds, after which the eyes are moved to the next location by means of a saccade (Henderson, 2003). Much modelling work has aimed to explain eye-gaze behaviour by visual saliency—that is, by low-level feature representations in the early visual cortex, by defining visual saliency as feature contrast between an image patch and its surround (Itti et al., 1998). Although these models can successfully pinpoint gaze-attracting locations in some scenes, they fall short when the scenes include attention-capturing higher-level features, such as faces (Cerf et al., 2009). Faces, and people in general, are especially effective in attracting the gaze (Yarbus, 1967; Birmingham et al., 2008), which likely reflects the central role of social cues for human behaviour and brain function (Hari et al., 2015). Moreover, the task given to the subject affects the gaze paths (Neider and Zelinsky, 2006; Torralba et al., 2006), and already the pioneering eye-tracking investigations by Buswell (1935) and Yarbus (1967) revealed that the fixations cluster on informative scene regions (for a review, see Henderson and Hollingworth, 1999).

The neural processes guiding gaze behaviour during natural vision have remained poorly understood. Moreover, brain activity associated with natural viewing can only be tracked with brain-imaging methods of high-enough temporal resolution. In this study, we used magnetoencephalography (MEG), which provides millisecond time resolution for noninvasive mapping of human cortical dynamics (for a review, see Hari and Salmelin, 2012). Our subjects viewed natural scenes, including landscapes and scenes depicting people in social interaction, while their eye gaze and MEG signals were monitored. We were especially interested in the initiation of the first saccade as an index of visual cues that likely are the most informative for scene understanding. By restricting our MEG analysis to the time-windows *preceding* the onset of the first saccade, we were able to avoid the contamination of the MEG data by oculomotor artifacts and to obtain information about brain processes that inform about the target of the first saccade.

A challenge to the analysis of MEG responses to even elementary (low-level) visual stimuli is that the cortical visual areas show substantial inter-individual variability in their size and position relative to anatomical landmarks, especially sulci (Amunts et al., 2000; Van Essen and Dierker, 2007), inducing great variability in the waveforms and spatial patterns of the evoked electromagnetic responses (Ales et al., 2010; Inverso et al., 2016). We therefore took a novel approach by applying representational similarity analysis (RSA; Kriegeskorte et al., 2008; Nili et al., 2014) to MEG data. That is, instead of assuming any specific shape for the waveform or topography of the MEG response, we made predictions on the similarity of the MEG-response patterns elicited by the scenes.

Figure 1 illustrates the analysis framework using simulated data. Random sources in the occipital poles (“early visual cortex”) had a correlation structure matching a categorical stimulus representation (Fig. 1A). The question is how this categorical response-geometry is reflected to the signals picked up by the MEG sensors. We constructed representational dissimilarity matrices (RDMs) by calculating the representational distance (1 – Pearson linear correlation) between each pair of the simulated MEG-response patterns from a set of neighbouring gradiometer sensors. The simulated-MEG-RDMs were compared with the ground-truth categorical-model-RDM using Spearman’s rank correlation (Fig. 1B). The analysis was repeated at each sensor location and the results were collected to a topographic map (Fig. 1C). The resulting maps were tested for significant correlation across subjects (Fig. 1D–E). As shown in Figure 1E, the occipital gradiometers correctly reflected the underlying categorical representation, thereby demonstrating the viability of the approach.

**Figure 1.**
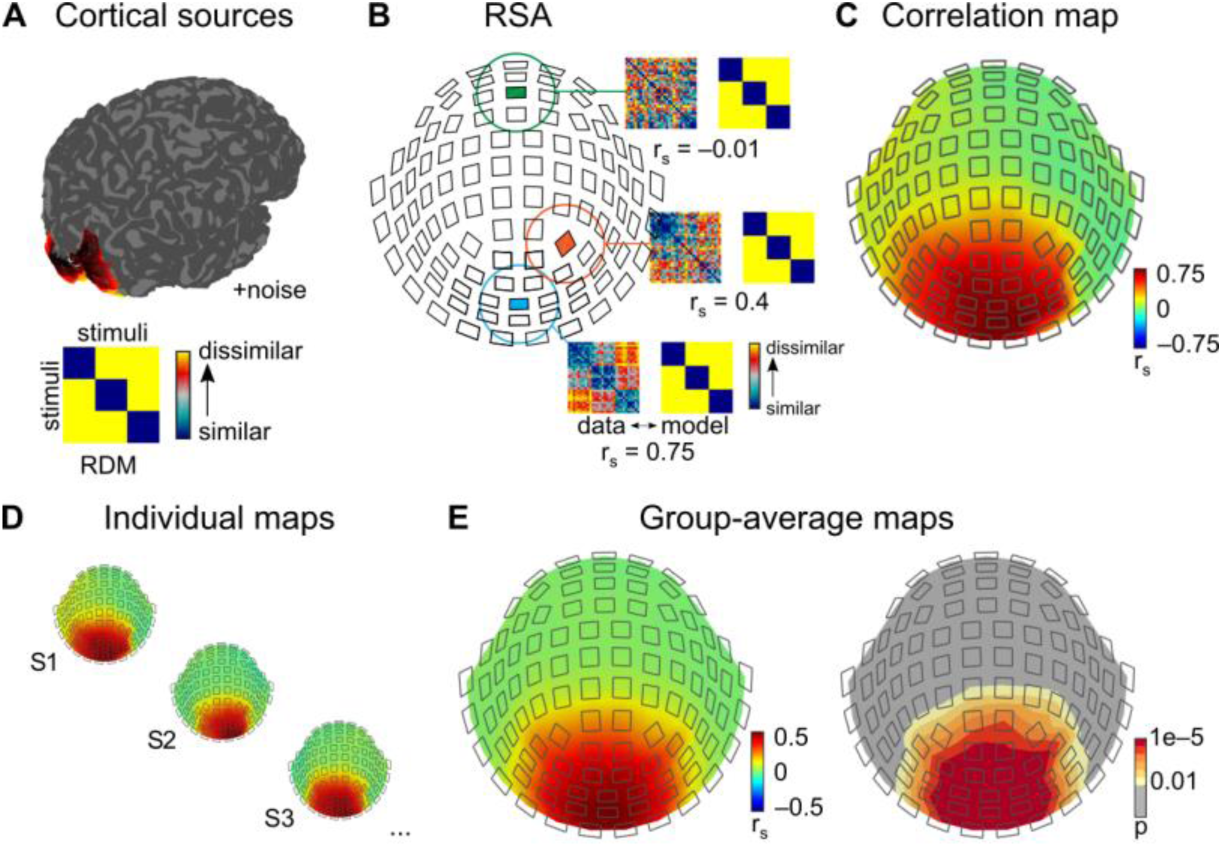
Representational similarity analysis for MEG data: a simulated categorical stimulus representation in early visual cortex. A) We demonstrate the idea of RSA for MEG using a simple simulation. We start by generating random patters in cortical nodes corresponding to the occipital poles in both hemispheres, with the correlation structure between the random patterns matching a categorical structure: 3 categories, 10 exemplars in each category (maximal correlation between exemplars of the same category, no correlation between exemplars of different categories). We calculate the forward solutions for the random patterns and add realistic noise to all patterns. B) The simulated responses are extracted in local neighbourhoods of MEG channels to construct an RDMs. The RDMs constructed from the simulated MEG data are compared to the ground-truth model-RMD (categorical model) using Spearman’s rank correlation. C) The Spearman’s correlation values are collected to a topographic map, and D) the individual topographic maps are E) averaged and tested for significant correlation (here simulated data from 10 subjects). The correlation is significant in occipital channels, matching the topography of the simulated sources.

In the present study, we looked for cortical signatures of the initiation of the first saccade. The representational distances between the MEG-response patterns elicited by the scenes were compared with predictions based either on the visual similarity of the stimuli or on the targets of the first saccades.

## MATERIALS AND METHODS

### Subjects

Eighteen healthy volunteers (10 males, 8 females; age range 18–39 yrs) with normal vision took part in this study. Ethics approval for the research had been obtained from the Ethics Committee of Hospital District of Helsinki and Uusimaa. Subjects gave written informed consent before participating in the study.

### Stimuli and experimental design

The stimuli were 199 photographs of natural scenes, including landscapes, scenes with single persons, and cluttered scenes with many people. The photographs were obtained from Wikimedia Commons™. The original photographs were cropped to size 1400 × 1050 pixels and converted to grayscale. The means and standard deviations of the grayscale values were matched across the stimulus set using the SHINE toolbox (Willenbockel et al., 2010). Figure 2 shows one stimulus image.

**Figure 2.**
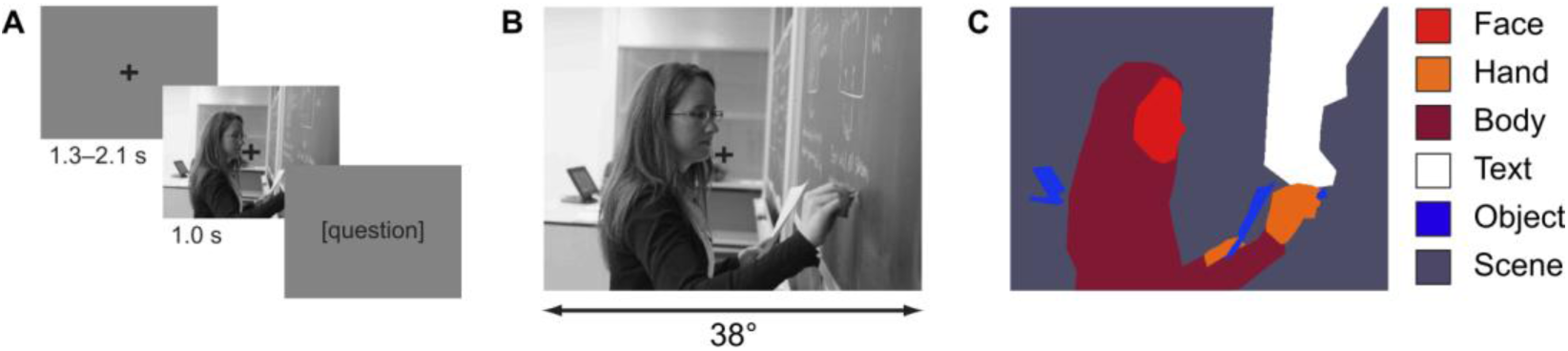
Natural scene stimuli and stimulus presentation. A) A fixation cross at the centre of the screen informed the subject about the onset of a new trial. The duration of the fixation screen was varied between 1.3 and 2.1 seconds to avoid subjects’ anticipating the onset of the stimuli. When the stimulus image appeared, the subject had been instructed to freely look at the image. The stimulus image was shown for 1.0 seconds, followed by a generic yes/no question about the content of the image. B) The stimuli were 199 photographs, converted to grayscale images. The diameter of a stimulus image was about 38 degrees. C) Each stimulus was manually annotated for features-of-interest: face, hand, body, text, object and background scene.

The stimuli were presented with a 3-DMD DLP projector (Panasonic ET-LAD7700/L) to a semitransparent screen at 105 cm viewing distance. The size of the stimulus image on the screen was 72 cm × 54 cm, giving a horizontal viewing angle of approximately 38 degrees. The timing of stimulus presentation (Fig. 2A) was controlled by PsychoPy (Peirce, 2007). Each trial begun with the fixation cross presented at the centre of the mid-gray screen for a random period of 1.3–2.1 s, followed by 1–0-s display of the stimulus image. The fixation cross was overlaid on the stimulus image but the subjects were instructed to freely move their eyes after the onset of the image. Stimulus offset was followed by a yes/no question in Finnish about the image (e.g., ‘Näkyikö kuvassa vettä?’ = ‘Was there water in the image?’; altogether 26 different questions). The questions were answered using an MEG-compatible response pad.

Altogether 199 different stimuli were shown in different random order for each subject. The images were divided to four experimental runs, separated with short breaks, with 50 of the 199 stimulus images presented in each run (49 in the last run). In addition, a set of 25 images, randomly selected of the 50 images of each run, was presented also upside-down in random order interleaved with the other stimuli. The data obtained for the inverted stimulus images are not analysed here.

The stimulus images were annotated for six features of interest: face, hand, body, text, object, and background scene (Fig. 2C). The annotations were manually drawn using the Matlab-based Object Labeling Tool from Derek Hoiem (http://dhoiem.cs.illinois.edu/software/).

### MEG data acquisition and pre-processing

MEG was recorded in a magnetically shielded room with a whole-scalp 306-channel MEG device (Elekta Oy, Helsinki, Finland) in the MEG Core, Aalto NeuroImaging, Aalto University, Finland. The device comprises 102 triple-sensor elements, with one magnetometer and two orthogonal planar gradiometers at each location. Only the gradiometer data (from 204 channels) were used in the analysis. The recording passband was 0.03–330 Hz, and the signals were sampled at 1000 Hz. The position of the subject’s head with respect to the MEG sensors was tracked throughout the experiment using four head position indicator (HPI) coils. In addition to the separate eye tracking (see below), horizontal and vertical electro-oculograms (EOGs) were recorded with the same recording passband and sampling rate as applied for the MEG.

The continuous MEG data were preprocessed using spatiotemporal signal-space separation (Taulu and Simola, 2006) implemented in MaxFilter software (Elekta Oy, Helsinki, Finland). This step included suppression of magnetic interference of external sources, compensation for head movement, and transformation of each individual’s data into a common head position. Single-trial MEG responses to stimulus images were extracted from the continuous MEG recording, baseline-corrected from –200 ms to 0 ms and low-pass filtered at 45 Hz using tools provided by the MNE and Fieldtrip software packages (Oostenveld et al., 2010; Gramfort et al., 2014).

For the representational similarity analysis, the signals were normalized by dividing the single-trial responses by the standard deviation of the response in a 200 ms time-window before the stimulus onset (baseline). Trials with blinks or poor fixation were rejected based on the eye-tracking data.

### Eye-gaze recording and data pre-processing

Eye gaze was tracked using an SR Research EyeLink1000 system (SR-Research Ltd., Ontario, Canada; sampling rate 500 Hz, average accuracy of gaze position better than 0.5 deg). The eye tracker was placed on a table in front of the subject. A 9-point calibration was performed at the start of the experiment and at the midpoint of the experiment. Fixations, saccades, and blinks were extracted from the continuous eye-tracking data during data collection using the software provided by the eye-tracker manufacturer. The extracted events were further processed in Matlab.

For each trial (stimulus), we extracted the onset of the first saccade and the position of the fixation before and after the first saccade. A trial was rejected if the fixation was not at the fixation cross at the onset of the trial, if the saccade amplitude was less than 0.5 deg, or if the subject had blinked. In addition, the data analysis was restricted to trials where the latency of the first saccade was between 125 ms and 350 ms. The eye-tracking data were drift-corrected based on the fixation locations at the onset of the trials.

### Representational-similarity analysis

The MEG responses to the different stimuli were compared with each other using correlation distance (1 – Pearson linear correlation). We applied a spatiotemporal searchlight approach, where the correlations were calculated between signals in overlapping sets of neighbouring MEG channels in time-windows of 25 ms (Fig. 3A, left panel). Channel neighbourhoods were defined using Fieldtrip (Oostenveld et al., 2010) and included on average 15 gradiometer pairs (channels at the edge of the MEG helmet, for example, had fewer neighbors than channels at the centre of the helmet). All pairwise comparisons between the stimuli were assembled in a representational dissimilarity matrix (RDM; Kriegeskorte et al., 2008; Nili et al., 2014) which, by definition, is symmetric and has a zero diagonal. The MEG-RDMs were calculated separately for each subject.

**Figure 3.**
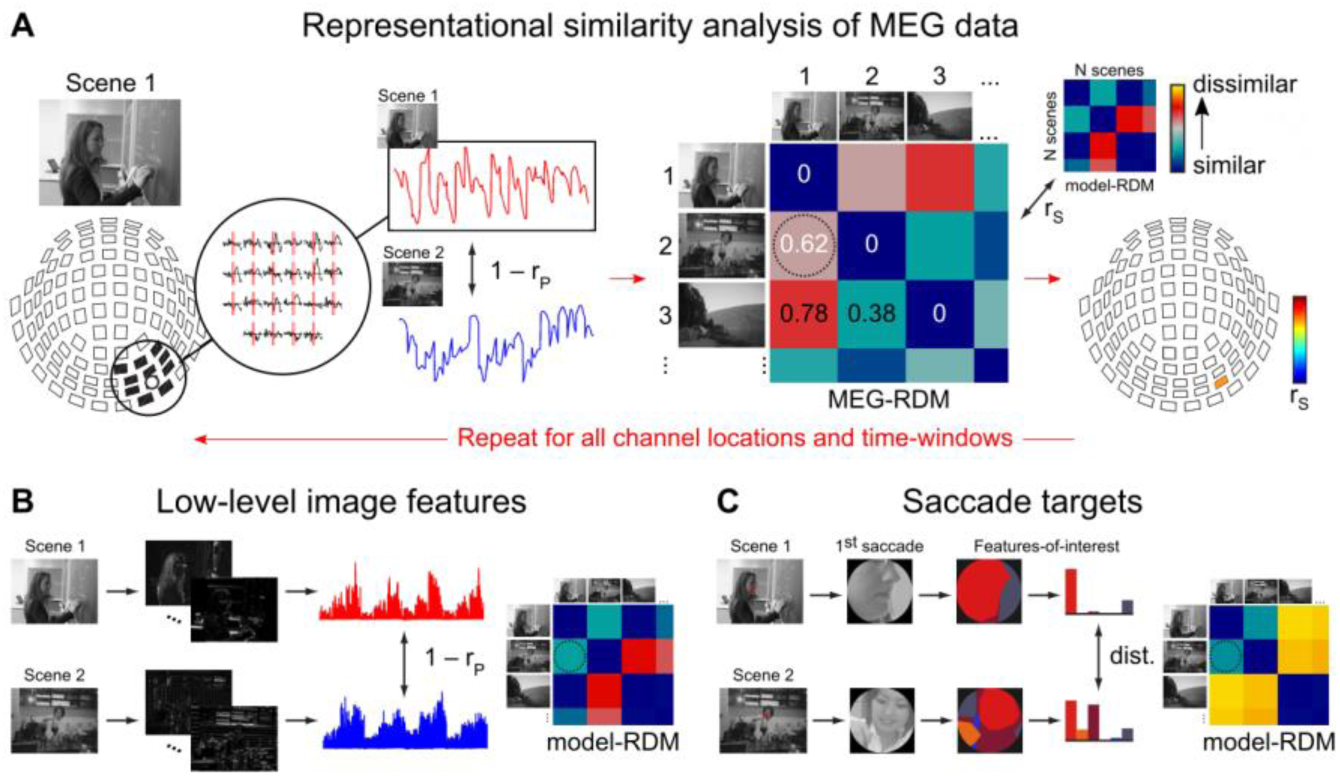
Representational similarity analysis. A) The MEG responses evoked by the different scenes were extracted in local spatiotemporal searchlights and compared to each other using correlation distance (1 – Pearson linear correlation). All pairwise comparisons between the MEG responses to the different scenes were collected to a representational dissimilarity matrix (RDM). The MEG-RDM was compared to a model-RDM using Spearman’s rank correlation (r_S_). The analysis was repeated for all channel locations (overlapping neighbourhoods) and for four non-overlapping time-windows. B) The stimulus images were convolved with Gabor wavelets to obtain a prediction of the low-level visual similarity of the stimuli. The convolution outputs were concatenated to have one feature vector for each image. All pairwise comparisons of the feature vectors were computed using correlation distance (r_P_) and the results were collected to an RDM. C) The saccade targets were characterized by extracting the features-of-interest (Fig. 2C from spotlights around the endpoint of the first saccade for each image. The similarity of saccade targets was estimated by calculating the Earth Mover’s Distance between the feature histograms.

The similarity of the RDMs across the group of subjects was studied to evaluate the amount of stimulus-related information in the RDMs, constructed from single-trial MEG responses. Using a leave-one-out approach, an RDM of one subject was compared with an RDM, where the RDMs of the other 17 subjects had been averaged. The comparison was based on Spearman’s rank correlation distance of the values in the upper triangle of the single-subject (left-out) RDM and the group-average (leave-one-out) RDM. The leave-one-out procedure was done for each subject, and the results were averaged and tested for significant deviation from zero. The analysis was repeated for all channel locations and in four time windows (25–50 ms, 50–75 ms, 75–100 ms, 100–125 ms).

Next, the MEG-RDMs were compared with different predictions on the representational similarity structure (model-RDMs): Gabor wavelet pyramid model (low-level visual similarity), similarity of saccade-target features, spatial distance between saccade end-point locations, and similarity of gaze scanpaths. In a model-RDM (Fig. 3B and 3C), each cell reflects the predicted dissimilarity of a stimulus pair. The native dimensions of an RDM were 199 × 199 (number of stimulus images). However, for each subject, some trials were excluded based on the eye-tracking data, and thus, the size of the RDMs differed between subjects. Note, however, that one benefit of the RSA approach is that exactly the same set of stimuli is not required for all subjects. The comparison between an MEG-RDM and a model-RDM was based on Spearman’s rank correlation distance of the values in the upper triangles of the RDMs (Kriegeskorte et al., 2008).

All comparisons were done on individual data and tested for statistically significant deviation from zero across the subjects. The multiple-comparisons problem was addressed by using a cluster-based permutation approach to test the statistical significance of the spatial cluster of significant sensors (sum of T-values; Maris and Oostenveld, 2007). The sensors showing statistically significant positive correlation (one-tailed t-test across subjects; p < 0.01) that survived the cluster-based permutation test (10 000 random permutations; p < 0.05) were visualized in topography plots.

#### Gabor-wavelet pyramid

A Gabor-wavelet pyramid model was adapted from the phase-congruency model provided by Kovesi (www.peterkovesi.com/matlabfns; Kovesi, 1999). Each stimulus image was represented with a set of Gabor wavelets of three spatial frequencies and four orientations at a regular grid over the image. The images were resized to 10% of the original size before the computation. The outputs of convolving the image with each of the Gabor wavelets were concatenated to have a representational vector for each image (Fig. 3B). The pairwise dissimilarities (1 – Pearson’s linear correlation) of these vectors were computed to obtain the Gabor-wavelet model-RDM for the stimuli.

#### Saccade targets

To characterize the target of the first saccade for each image in each subject, we defined a spotlight with a diameter of 5.2 deg (about the size of fovea; Wandell, 1995) to extract the features around the landing position of a saccade. The features were extracted from the annotated stimulus images (Fig. 2C). The similarity of the feature histograms between two targets was estimated using Earth Mover’s Distance. The distances were collected to an RDM (Fig. 3C). Because the RDMs were based on individual eye-tracking data, we had a unique reference RDM for each subject based on their saccade targets.

#### Saccade landing positions

The distances between saccade landing positions between each pair of stimulus images were calculated from saccade vectors, that is vectors from the start point of the saccade (fixation at stimulus onset) to the endpoint of the first saccade. The distances were collected to an RDM, separately for each subject.

#### Scanpaths

The full gaze scanpath for each image was characterized using the ScanMatch toolbox (Cristino et al., 2010). Each stimulus image was divided into 11 bin × 9 bin and the temporal binning was set to 50 ms. The similarity of two scanpaths was quantified using the Needleman-Wunsch sequence alignment algorithm implemented in the ScanMatch toolbox (Cristino et al., 2010). The scanpath similarities between all pairs of images were computed and collected to an RDM. The scanpath similarities were characterized separately for each subject.

## RESULTS

### The first saccade is typically directed to people—especially to their faces

Figure 4 shows the landing positions of the first saccades in all stimulus images, with one point for each individual subject. The gaze was typically drawn to faces or bodies of the persons in the scene. If the scene contained no persons, a salient object (such as a football or a boat) often attracted the first saccade. Although the landing positions of the first saccades were largely consistent across the subjects, for some scenes the first saccade clearly diverged between individuals; see, for example, the scene depicting a woman putting a book on a shelf (Fig. 4, seventh row, last image), where the viewers’ gaze was drawn either to the woman’s face or to the acting hand. Hence, we used the individual instead of group-average results for the analysis where we related the eye-gaze and MEG data.

**Figure 4.**
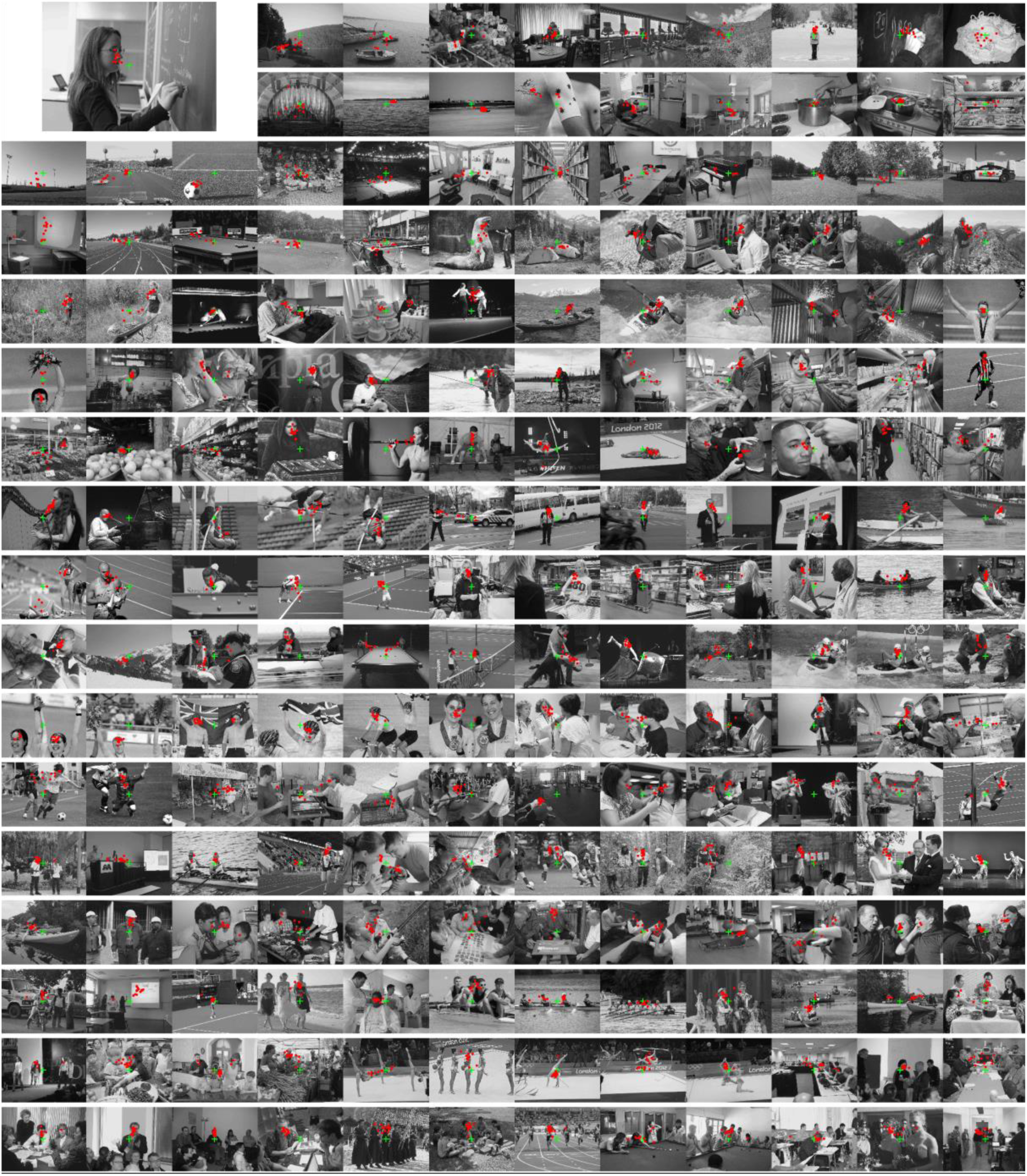
Natural scene stimuli overlaid with the eye-gaze results. Subjects were instructed to look freely at the scenes. The red dots indicate the locations of the first saccade for indivudual subjects. The fixation cross is shown as a green cross (a black fixation cross was used in the experiments, Fig. 2A). Note the clustering of the first saccades to similar targets across the group of subjects.

To characterize the saccade targets in each image, we defined a gaze-spotlight around the landing position of each saccade to match the size of the fovea in the visual field (diameter 5.2 deg). Figure 5 shows the group-average gaze-spotlights, separately for each of the 199 scenes. The midpoint of each group-average gaze-spotlight was defined from the individual eye-gaze data by convolving the individual saccade landing positions with a Gaussian kernel and finding the peak of the resulting heatmap. The group-average spotlights are shown to visualize the most common features in the scenes that attracted the gaze. Figure 6 shows the same gaze-spotlights with annotated features-of-interests; the predominance of the red color indicates that most of the saccade targets contained faces, and a minor part objects (blue color). The annotations were based on previous studies suggesting that faces, people, and text attract the gaze (Yarbus, 1967; Cerf et al., 2009).

**Figure 5.**
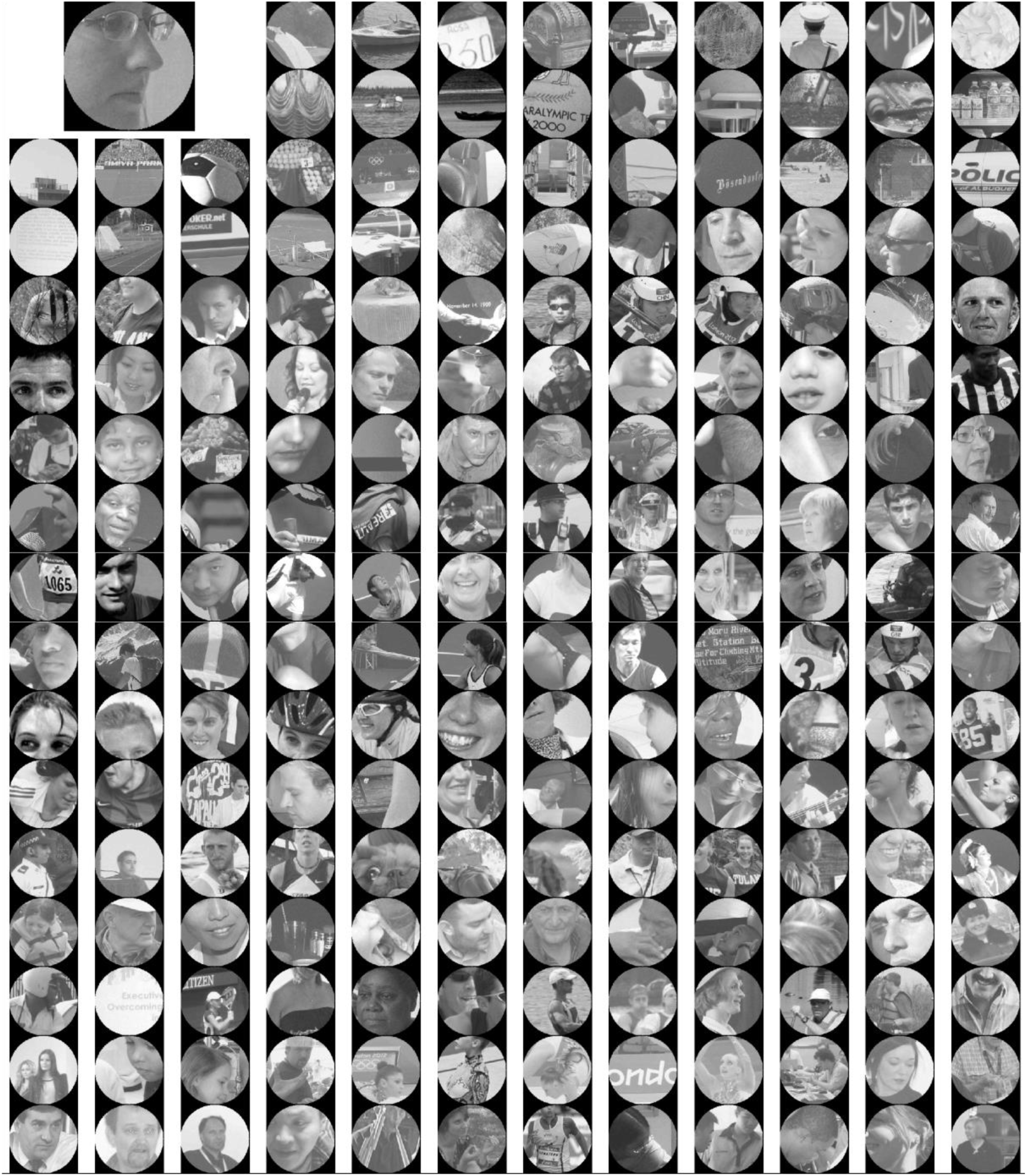
Average targets of the first saccades. For each scene, we defined a spotlight around the endpoint of the first saccade. The size of the spotlight was about the size of the fovea in the visual field (diameter 5.2 degrees). Here the group-average spotlights are shown for all 199 scenes.

**Figure 6.**
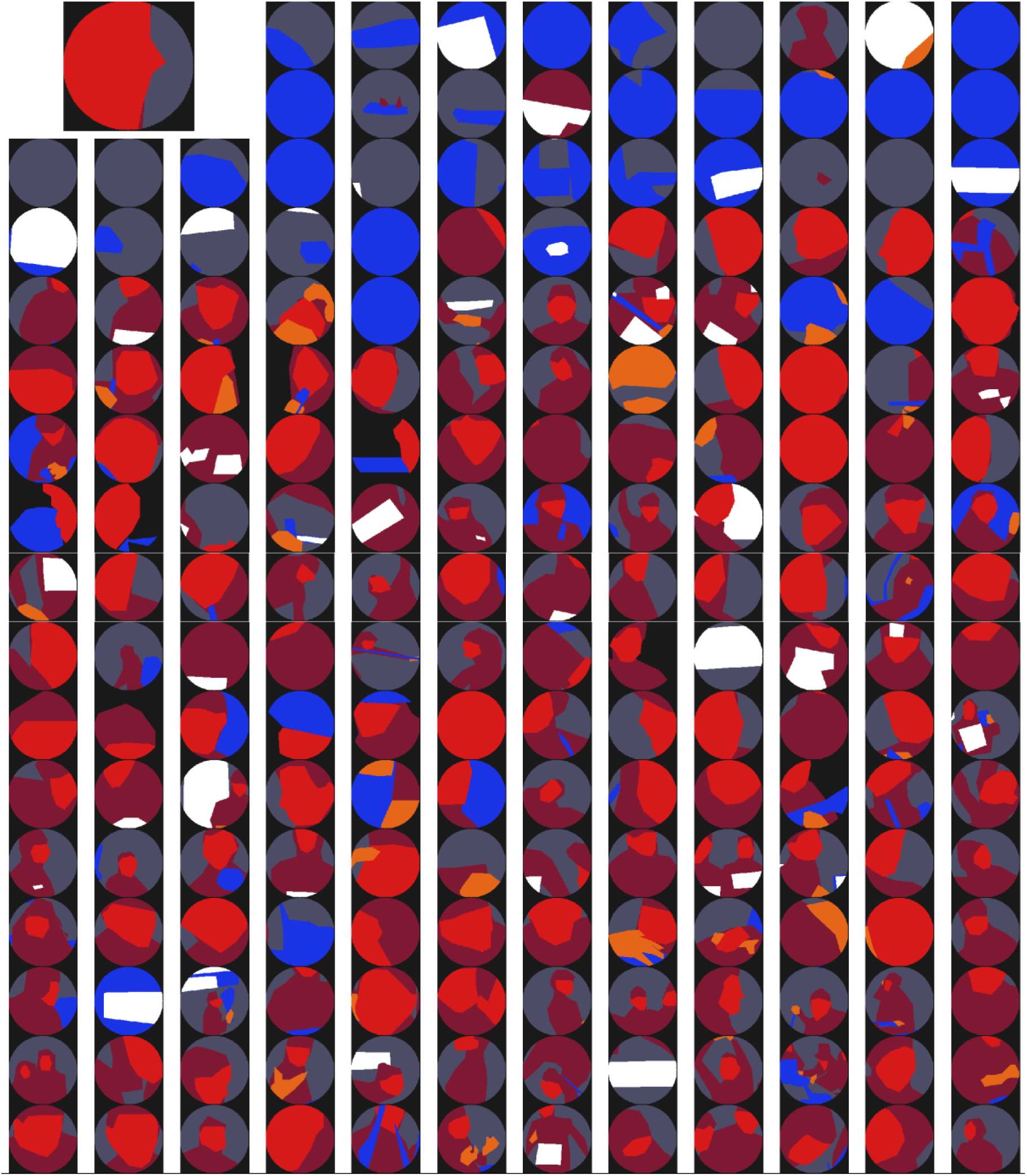
Annotations of the saccade targets. To quantify the features within the saccade targets, all images were annotated for features-of-interest (Fig. 2C). Here the annotated group-average saccade targets are shown for all 199 scenes (red = face, orange = hand, dar red = body, white = text, blue = object, dark blue = background scene).

The features-of-interest were quantified from the gaze-spotlights around the landing positions of the first saccades, separately for each subject. Figure 7 shows that the amounts of features-of-interests were greater in the saccade spotlights (coloured bars) than in the whole stimulus images (gray bars), except for the pixels depicting background scene which were less pronounced within the saccade spotlights. The proportional increases of the feature-of-interest pixels in the gaze spotlights compared with the whole images were greatest for faces and text, which only occupied a small portion of the stimulus images but large proportions of the gaze-spotlights of the first saccades.

**Figure 7.**
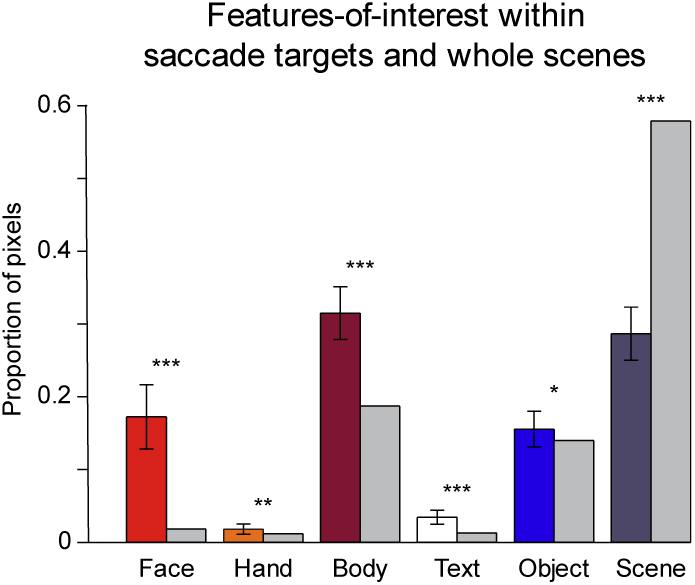
Features-of-interest within saccade targets and whole stimulus images. Results on the proportion of different features-of-interest (see also Figs. 2C and 6) within the saccade targets and in the whole stimulus images are shown. Each coloured bar indicates the mean proportion of each of the features-of-interest (face, hand, body, text, object, background scene) within the spotlights of the first saccades. The error bars indicate the standard deviation across the 18 subjects. The gray bars indicate the proportion of each of the features-of-interest in the whole stimulus image. The asterisks indicate statistically significant difference (two-tailed t-test; ***p<0.001, **p<0.01, *p<0.05).

### Single-trial MEG responses carry information about natural-scene stimuli

While the subjects were viewing the scenes, we followed their brain dynamics using MEG. The dynamic cortical representations were characterized by means of representational distance matrices constructed from the single-trial MEG data (for an overview of the analysis, see Fig. 3A). To construct an MEG-RDM, we extracted signals from a set of neighbouring gradiometers in a 25-ms time-window and calculated all pair-wise correlations between the signals elicited by the different scenes. The MEG-RDMs were constructed separately for each channel location and in four different time-windows. The RDMs can be directly compared between different individuals (without any need for matching the MEG response shapes between individuals), or an MEG-RDM can be directly compared with an RDM describing the representational geometry of a computational model (Kriegeskorte et al., 2008; Kriegeskorte and Kievit, 2013).

First to confirm the suitability of the single-trial MEG data for representational similarity analysis, we studied the similarity of the RDMs across the group of subjects (Fig. 8A). We would hypothesize that all stimulus images do not evoke equally distinct responses but that some stimuli evoke more similar responses, and thus, the ranking from the least to the most similar stimulus pair would be correlated across subjects. The RDM of a single subject was compared to the group-average RDM of the other subjects by calculating the Spearman’s rank correlation between the RDMs. To avoid contamination of the MEG signals by oculomotor activity and to learn about brain activity supporting the selection of the first saccade target, we restricted the analysis to the part of the MEG response before the onset of the first saccade. RDMs showed similarity across the group of subjects starting from the time-window of 50–75 ms, first in the occipital cortex (Fig. 8A). That is, the MEG-RDMs showed replicable structure across the group of subjects from the onset of the stimulus-related response. At later time-windows, the subjects’ RDMs are continuously correlated, with peaks in the occipital cortex but extending also to temporal and parietal regions, with right-hemisphere dominance.

**Figure 8.**
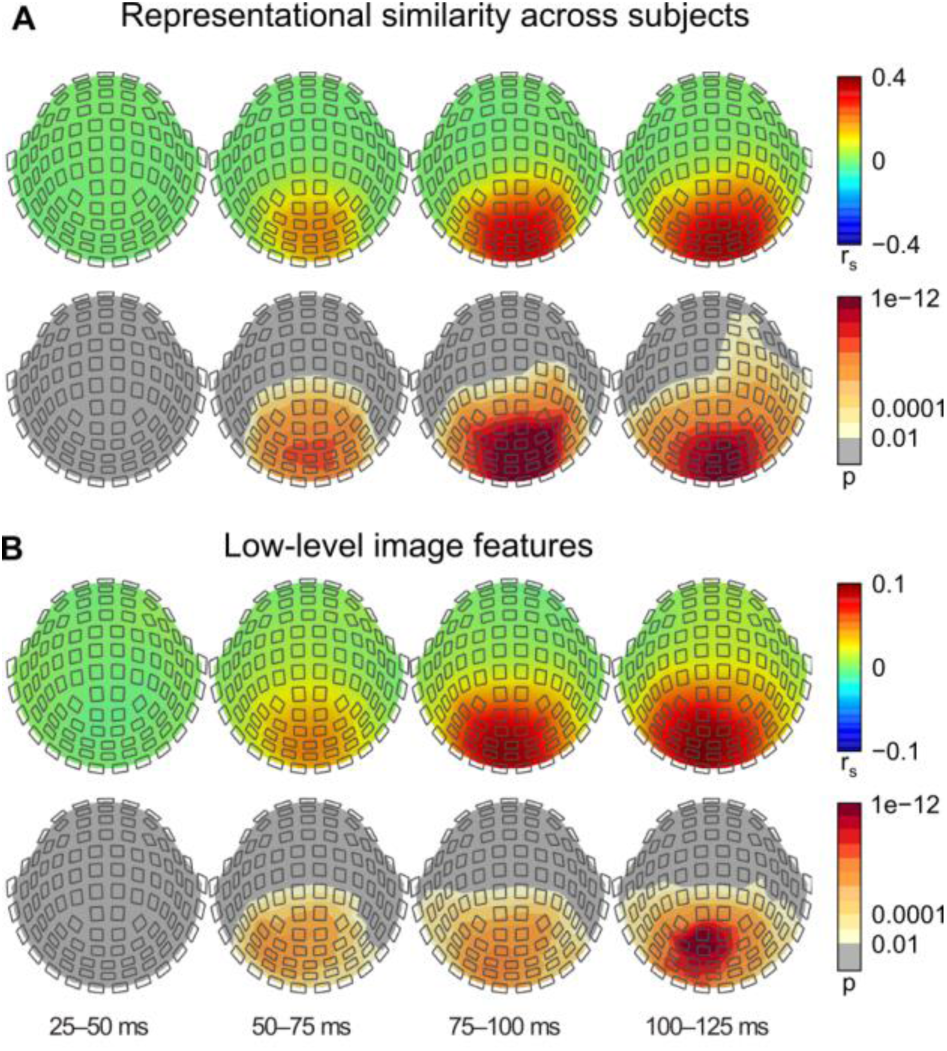
Representational similarity of single-trial MEG-RDMs across subjects and comparison with low-level image features. A) The upper row shows the mean Spearman’s rank correlation between the RDM of a leave-one-out-subject and the average RDM of the other 17 subjects, calculated separately within a spatiotemporal searchlight at each channel location. The analysis was done separately in four time-windows: 25–50 ms, 50–75 ms, 75–100 ms, and 100–125 ms. The bottom row shows the corresponding p-values (one-tailed t-test, n = 18; the significance of a spatial cluster tested by permutation, p < 0.05). The MEG-RDMs showed similar structure across subjects from the time-window 50–75 ms from stimulus onset, first above the occipital cortex. B) The upper row shows the mean Spearman’s rank correlation between the RDM constructed based on the low-level visual similarity of the stimulus images (Fig. 3B) and the MEG-RDMs, calculated separately within a spatiotemporal searchlight at each channel location and in four time-windows: 25–50 ms, 50–75 ms, 75–100 ms, and 100– 125 ms. The bottom row shows the corresponding p-values (one-tailed t-test, n = 18; the significance of a spatial cluster tested by permutation, p < 0.05). The low-level visual similarity of the stimuli explained structure in the MEG-RDMs from the time-window 50– 75 ms onwards.

### Similarity of low-level image features explains similarity of early MEG responses

The Gabor wavelet model is commonly considered as the standard model of visual representations of area V1 (Carandini et al., 2005), and has been shown to explain stimulus representations in areas V1–3 when characterized using functional magnetic resonance imaging (Kay et al., 2008; Henriksson et al., 2015). Figure 8B shows the results when the MEG-RDMs were compared with an RDM constructed based on the Gabor wavelet pyramid model of the stimuli, reflecting the representation of low-level image features. The comparison was based on the rank correlation between model-RDM and the MEG-RDMs. The results in Figure 8B resemble the results shown in Figure 8A, where we assessed the similarity of the RDMs across the group of subjects. Thus, from the early time-window (50–75 ms), the similarity of the low-level image features of the stimuli explained similarity of MEG responses (Fig. 8B).

### Saccade targets explain MEG response-pattern similarity at 75–100 ms

Our main objective was to combine MEG and eye-tracking data to uncover cortical dynamics of saccade-target selection during free-viewing of scenes. The eye-tracking results revealed the features in the stimulus images that captured the subjects’ attention. The features-of-interest were extracted from the annotated images for the first saccades, separately for each subject. The similarity of each pair of feature histograms (number of pixels for each feature-of-interest within a saccade spotlight) was estimated and collected to an RDM (for an example, see Fig. 3C). The RDMs were constructed separately for each subject based on the individual eye tracking results and compared with the MEG-RDMs. Figure 9A shows that the similarity of visual features within the saccade targets explained the similarity of the MEG responses starting 75–100 ms after stimulus onset, first in the right-hemisphere parieto-occipital cortex (Fig. 9A).

**Figure 9.**
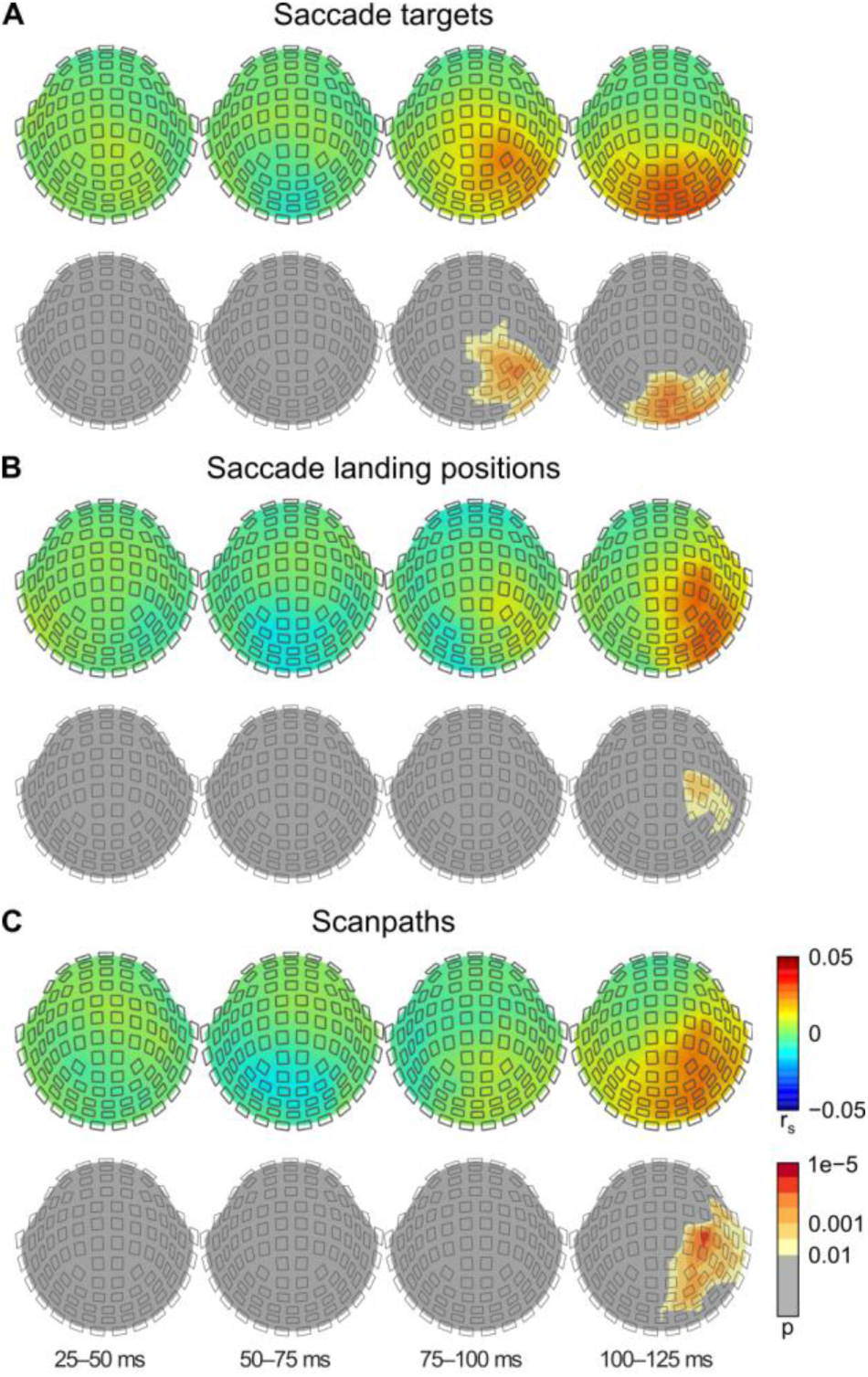
Comparisons of MEG-RDMs with targets and locations of the upcoming saccades. A) The saccade target similarity between two stimuli was characterized from the features-of-interest within the saccade spotlights (see also Figs. 2C, 3C and 6). The upper row shows the mean Spearman’s rank correlation between the RDMs constructed based on the similarity of the features within the individual saccade targets and the individual MEG-RDMs, calculated separately within a spatiotemporal searchlight at each channel location and in four time-windows: 25–50 ms, 50–75 ms, 75–100 ms, and 100– 125 ms. The bottom row shows the corresponding p-values (one-tailed t-test, n = 18; the significance of a spatial cluster tested by permutation, p < 0.05). The similarity of the saccade targets explained structure in the MEG-RDMs from the time-window 75–100 ms onwards. B) RDMs were constructed based on the distance between saccade landing positions, and correlated with the MEG-RDMs. Upper row shows the mean Spearman’s rank correlation and bottom row the corresponding p-values. C) RDMs were constructed based on the similarity of the full scanpaths of the eye movements between the different scenes, and correlated with the MEG-RDMs. Upper row shows the mean Spearman’s rank correlation and bottom row the corresponding p-values. Scanpath similarity explained structure in the MEG-RDMs first in time-window 100–125 ms after stimulus onset, in right-lateralized sensors.

Next, we built an RDM for each subject based on the landing positions of the first saccades. The MEG-RDMs from a cluster of right-lateralized sensors correlated with the saccade-location RDMs in the time-window 100–125 ms after stimulus onset (Fig. 9B). Finally, we constructed RDMs based on, not only on the first saccade, but on the similarity of the full scanpaths of eye movements for each scene. The similarity between each pair of eye-movement sequences was quantified using the ScanMatch toolbox (Cristino et al., 2010). The scanpath similarity explained similarity in the MEG responses in the time-window 100–125 ms after stimulus onset, over right-lateralized sensors (Fig. 9C).

### Stimulus vs. saccade-aligned analysis

Thus far, results are presented from analyses of MEG signals time-locked to the stimulus onset, thus ignoring variability in saccade latencies. Can we see any effects of saccade latency in the MEG signals? Figure 10A shows grand-average MEG responses from occipital (top panels), right temporal (middle panels) and frontal (bottom panels) channels (root-mean-square amplitudes of signals in gradiometer pairs averaged across the channels and subjects) time-locked both to the stimulus onset (left column) and to the saccade onset (right column). The responses were binned to eight equal-sized groups based on saccade latency (shown in different colours). Responses time-locked to the stimulus onset show no clear evidence for saccade-latency-related effects on amplitude or latency before the onset of the saccade. Similarly, when the responses were time-locked to the onset of the saccade, the post-saccadic responses show no clear saccade-latency-related effects (Fig 10A, right column). Figure 10B shows also the grand-average EOG responses (root-mean-square amplitudes of signals in vertical EOG pair averaged across subjects) to compare with the MEG responses, and to evaluate whether the oculomotor artefact interferes with the interpretation of the MEG response.

**Figure 10.**
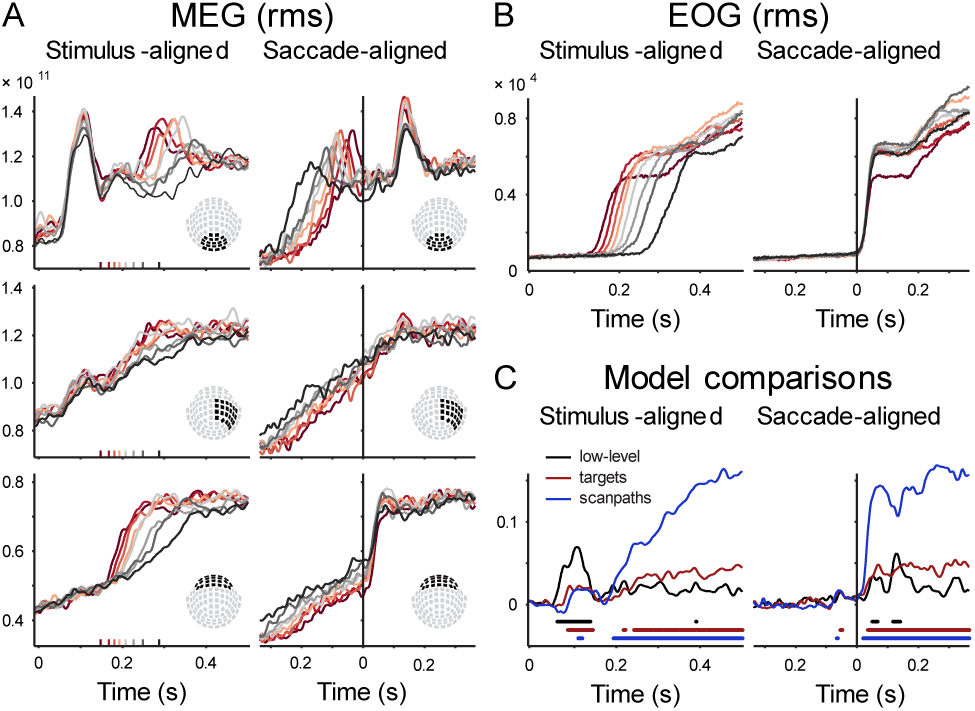
Stimulus vs. saccade-aligned analysis. A) Grand-average MEG responses from occipital (top-row), right temporal (middle row) and frontal channels (bottom-row) time-locked both to the stimulus onset (left column) and to the saccade onset (right column) are shown. The grand-average responses were calculated as the root-mean-square amplitudes of signals in gradiometer pairs averaged across the channels and subjects. The responses were binned to eight equal-sized groups based on saccade latency (shown in different colours). B) Grand-average EOG signals are shown aligned to stimulus onset (left column) and saccade onset (right column). The signals were binned to eight categories based on the saccade latency before averaging (shown in different colours). C) An MEG-RDM was constructed from all gradiometer signals, separately for each time-point, and compared with model-RDMs: low-level visual similarity of the stimulus images (shown in black), saccade target similarity (shown in red), and scanpath similarity (shown in blue). The results are shown as Spearman’s rank correlation between the MEG-RDM and model-RDMs averaged across subjects (lines indicate significant correlation; one-tailed t-test, FDR < 0.01).

We also performed the RSA analysis on the MEG responses time-locked to the saccade-onsets instead of stimulus-onset. Figure 10C shows the time-resolved results on the comparisons between the MEG-RDMs and the model-RDMs, both time-locked to the stimulus (left column) and saccade onset (right column). The MEG-RDMs were constructed from signals from all gradiometers, separately for each time-point. When time-locked to the stimulus onset, the MEG-RDMs and model-RDMs in the pre-saccadic time-window (< 125 ms) exhibit the same behaviour as shown in Figures 8–9. Moreover, the sustained post-saccade (> 200 ms) correlation between the MEG-RDMs and the saccade-target-RDMs (red curve) suggests sustained information about the saccade targets in the brain activity. The comparison of the MEG-RDMs with the scanpath-RDM (blue curve), on the other hand, is likely contaminated by oculomotor (and not brain) signals in the post-saccadic time-window. When time-locked to the saccade onsets (Fig. 10C, right panel), the MEG signals show little evidence for any saccade-related effects before the saccade onset. After the saccade onset, oculomotor signals likely explain the high correlation between the MEG-RDMs and scanpath-RDMs. On the other hand, the sustained correlation between the MEG-RDMs and the saccade-target-RDMs (red curve) suggests sustained information about the identity of the saccade targets in the post-saccadic brain responses. In addition, the peak in the correlation with the low-level visual features coincides with the peak in the MEG responses from the occipital sensors (Fig. 10A, top row, right column). Taken together, signatures of saccade planning are evident in the MEG responses aligned to the stimulus-onset (Fig. 9 and Fig. 10, left column) but not in the responses aligned to the saccade-onset (Fig. 10, right column).

## DISCUSSION

What are the neural mechanisms underlying guidance of saccades during natural visual behaviour? What attracts our gaze in the absence of explicit instructions what to look at? In this study, we presented natural visual scenes to healthy humans whose brain responses and eye gaze were simultaneously tracked. Although the subjects were not given instruction on where to look at in the images, they were remarkably consistent in where they made the first saccade in each scene. Moreover, we were able to relate the evoked single-trial MEG responses to the natural-scene stimuli, starting at time-window 50–75 ms, and more interestingly, to the target of the upcoming first saccade at 75–100 ms from stimulus onset, in other words, before the saccade onset.

Combination of eye-tracking and MEG data enabled us to follow the cortical dynamics leading to saccade initiation, and thus, to address questions on saccade-target selection during natural visual behaviour. MEG with its millisecond time-resolution is better suited to be combined with eye tracking than is functional magnetic resonance imaging (fMRI), which is inherently limited in its temporal resolution by the sluggish hemodynamic response. To relate the complex MEG responses to the eye-tracking results, we applied representational similarity analysis. RSA of fMRI data has proven powerful in unraveling visual activation patterns across the hierarchy of visual processing pathways and in relating cortical activations with computational models and behaviour (Kriegeskorte et al., 2008; Kriegeskorte and Kievit, 2013; Khaligh-Razavi and Kriegeskorte, 2014). Instead, only few previous studies have used RSA together with MEG data (see, *e.g.*, Carlson et al., 2013; Cichy et al., 2014), and ours is the first study to apply RSA to combined eye-gaze and MEG data. Another novel feature in our analysis is that instead of using all MEG channels, we performed the RSA analysis in local neighbourhoods of gradiometer channels, hence preserving some spatial information in the resulting topographic maps. This approach was inspired by the searchlight approach widely used together with fMRI data, where the whole brain-volume is scanned using a local searchlight at each location in the brain to perform multivariate analysis on the local brain-activity patterns (Kriegeskorte et al., 2006). Our results on the early visual feature-representations in the occipital cortex match well with previous fMRI findings on low-level feature representations in visual areas V1–3 (Kay et al., 2008; Henriksson et al., 2015). Furthermore, our right-hemisphere-lateralized results on saccade-planning are consistent with activation in fronto-parietal brain regions, predominantly in the right hemisphere during stimulus-driven attention (Corbetta and Shulman, 2002). Fom the channel-level analysis of the MEG responses, we are, however, unable to make precise conclusions about the underlying cortical sources. A promising approach for future studies is to combine MEG with fMRI data though the RSA framework (Cichy et al., 2016).

We restricted our MEG analysis to the early responses (up to 125 ms) leading to saccade initiation; in only a few cases was the latency of the first saccade shorter than 125 ms, and these traces were rejected from the analysis. An intriguing question for future research will be the neural origins of the within- and between-individual variability in saccade latencies during natural gaze behaviour. Our results that the brain signatures of saccade planning are evident in MEG traces aligned to stimulus onset, but not in traces aligned to saccade onset, support previous findings suggesting that saccade-latency variability is related to motor planning of the saccades instead of perceptual processes based on sensory information that starts to accumulate immediately after the scene onset (Thompson et al., 1996). However, some of the saccade-latency variability could also be related to the content of the stimulus scenes, especially to the eccentricity of the saccade-target, and thus to cortical processing of the obtained sensory information.

Visual acuity, and therefore the effective sampling of visual information, is by far the highest in the central retina, which is the very reason why saccades are made during natural visual exploration. Ensemble representations have recently been suggested to explain our subjectively rich percept of the visual world and also to provide a foundation for guiding eye gaze (Cohen et al., 2016). What can we then perceive with the low-acuity peripheral vision to guide our saccades? The present results, together with previous findings (Henriksson and Hari, in press; Birmingham et al., 2008; Fletcher-Watson et al., 2008; Crouzet et al., 2010), suggest that social cues, such as faces and people, capture the attention and eye-gaze even in the peripheral visual field. Using a saccadic forced-choice task, a recent study demonstrated that faces can be categorized even in the far-periphery (up-to 80 deg), with an advantage in terms of categorisation speed in comparison to other target categories, including animals and vehicles (Boucart et al., 2016). A saccade toward a face can even be difficult to suppress when the task has been to look at something else (Crouzet et al., 2010). Overall, eye movements in natural behaviour are dominated by contextual and high-level features, including social cues (Henriksson and Hari, in press; Cerf et al., 2009), instead of low-level visual saliency (Itti and Koch, 2001).

Given that our stimuli were natural scenes, they were shown only for one second, and that other objects within the scene might have been more informative when answering the generic question asked after the stimulus, it could be considered surprising how consistent our subjects were when making the first saccade toward the people within the scenes. Instead of merely confirming the attention-capturing power of the faces, our result suggest that the contents of a scene are best understood by observing the actors within the scene. That is, whereas the rapid gist of a scene might be perceived without any analysis of the individual objects within the scene (Oliva and Torralba, 2006; Cohen et al., 2016), the more detailed understanding of the scene might be provided not from the objects but from the people acting within the scene. This consideration is consistent with the recent finding that human scene categorization is better explained by the function of a scene, such as potential for shopping or hiking, than by the individual objects within the scene (Greene et al., 2016). Furthermore, semantic consistency between a foreground object and a background scene can facilitate the perception of both the object and the scene (Davenport and Potter, 2004). Although some of the object–scene pairs in the study by Davenport and Potter (2004) were actor–scene pairs (*e.g.*, a priest/footballer and a church/football field), the study did not address any possible difference between the actor– scene and object–scene pairs, thus leaving open the question of the superiority effect of consistent actor–scene pairs as compared with object–scene pairs for scene understanding. Recent modelling work has also been motivated by the idea that modelling the people within the scene facilitates scene understanding. The extraction of the 3D layout of a scene can be improved by identifying also the people within the scene (Fouhey et al., 2014). To interpret the behaviour of the persons within a scene might require an approach that follows the gaze of the persons (Recasens et al., 2015). Whereas humans are naturally cued to follow others’ gaze and to orient attention to the same object that the others are looking at (joint attention), this is not an easy task for computer vision.

In conclusion, we have introduced a novel approach for combining MEG and eye-tracking data and demonstrated that single-trial MEG responses can be related to the upcoming saccades during free-viewing of natural scenes. We believe that the initiation of the first saccade is informative about the rapid cortical processing leading to scene understanding, and that the proposed data-analysis approach can be applied to address various other questions related to natural visual behaviour. Moreover, the finding that the viewers automatically directed their gaze towards any people in a scene leads to ask whether an actor within a scene would provide the best cues for scene understanding.

## Acknowledgements

We thank Mia Illman for help in MEG measurements and Veli-Matti Saarinen for help in eye-tracking measurements. The annotations of the stimulus images were created using the Object Labeling Tool from Derek Hoiem, University of Illinois. This work was financially supported by the Academy of Finland Postdoctoral Researcher Grant (278957; LH), the Louis-Jeantet Foundation (RH), and Strategic Centre for Science, Technology and Innovation in Health and Well-being (SHOK, SalWe, Finland; RH). The authors declare no competing financial interests.

## REFERENCES

Ales J, Carney T, Klein SA (2010) The folding fingerprint of visual cortex reveals the timing of human V1 and V2. Neuroimage 49:2494–2502.

Amunts K, Malikovic A, Mohlberg H, Schormann T, Zilles K (2000) Brodmann’s areas 17 and 18 brought into stereotaxic space—where and how variable? Neuroimage 11: 66–84.

Birmingham E, Bischof WF, Kingstone A (2008) Social attention and real-world scenes: The roles of action, competition and social content. Q J Exp Psychol 61:986–998.

Boucart M, Lenoble Q, Quettelart J, Szaffarczyk S, Despretz P, Thorpe SJ (2016) Finding faces, animals, and vehicles in far peripheral vision. J Vis 16:1–13.

Buswell GT (1935) How people look at pictures. University of Chicago Press Chicago.

Carandini M, Demb JB, Mante V, Tolhurst DJ, Dan Y, Olshausen BA, Gallant JL, Rust NC (2005) Do we know what the early visual system does? J Neurosci 25:10577–10597.

Carlson T, Tovar DA, Alink A, Kriegeskorte N (2013) Representational dynamics of object vision: the first 1000 ms. J Vis 13:1–1.

Cerf M, Frady EP, Koch C (2009) Faces and text attract gaze independent of the task: Experimental data and computer model. J Vis 9:10–10.

Cichy RM, Pantazis D, Oliva A (2014) Resolving human object recognition in space and time. Nat Neurosci 17:455–462.

Cichy RM, Pantazis D, Oliva A (2016) Similarity-based fusion of MEG and fMRI reveals spatio-temporal dynamics in human cortex during visual object recognition. Cereb Cortex:bhw135.

Cohen MA, Dennett DC, Kanwisher N (2016) What is the bandwidth of perceptual experience? Trends Cogn Sci 20:324–335.

Corbetta M, Shulman GL (2002) Control of goal-directed and stimulus-driven attention in the brain. Nat Rev Neurosci 3:201–215.

Cristino F, Mathôt S, Theeuwes J, Gilchrist ID (2010) ScanMatch: A novel method for comparing fixation sequences. Behav Res Methods 42:692–700.

Crouzet SM, Kirchner H, Thorpe SJ (2010) Fast saccades toward faces: face detection in just 100 ms. J Vis 10:1–17.

Davenport JL, Potter MC (2004) Scene consistency in object and background perception. Psychol Sci 15:559–564.

Fletcher-Watson S, Findlay JM, Leekam SR, Benson V (2008) Rapid detection of person information in a naturalistic scene. Perception 37:571–583.

Fouhey DF, Delaitre V, Gupta A, Efros AA, Laptev I, Sivic J (2014) People watching: Human actions as a cue for single view geometry. Int J Comput Vis 110:259–274.

Gramfort A, Luessi M, Larson E, Engemann DA, Strohmeier D, Brodbeck C, Parkkonen L, Hämäläinen MS (2014) MNE software for processing MEG and EEG data. Neuroimage 86:446–460.

Greene MR, Baldassano C, Esteva A, Beck DM, Fei-Fei L (2016) Visual scenes are categorized by function. J Exp Psychol Gen 145:82.

Hari R, Henriksson L, Malinen S, Parkkonen L (2015) Centrality of social interaction in human brain function. Neuron 88:181–193.

Hari R, Salmelin R (2012) Magnetoencephalography: from SQUIDs to neuroscience: Neuroimage 20th anniversary special edition. Neuroimage 61:386–396.

Henderson JM (2003) Human gaze control during real-world scene perception. Trends Cogn Sci 7:498–504.

Henderson JM, Hollingworth A (1999) High-level scene perception. Annu Rev Psychol 50:243–271.

Henriksson L, Hari R (in press) Contextual and social cues may dominate natural visual search. Behav Brain Sci.

Henriksson L, Khaligh-Razavi S-M, Kay K, Kriegeskorte N (2015) Visual representations are dominated by intrinsic fluctuations correlated between areas. NeuroImage 114:275–286.

Inverso SA, Goh X-L, Henriksson L, Vanni S, James AC (2016) From evoked potentials to cortical currents: Resolving V1 and V2 components using retinotopy constrained source estimation without fMRI. Hum Brain Mapp 37:1696–1709.

Itti L, Koch C (2001) Computational modelling of visual attention. Nat Rev Neurosci 2:194–203.

Itti L, Koch C, Niebur E (1998) A model of saliency-based visual attention for rapid scene analysis. IEEE Trans Pattern Anal Mach Intell 20:1254–1259.

Kay KN, Naselaris T, Prenger RJ, Gallant JL (2008) Identifying natural images from human brain activity. Nature 452:352–355.

Khaligh-Razavi S-M, Kriegeskorte N (2014) Deep supervised, but not unsupervised, models may explain IT cortical representation. PLoS Comput Biol 10:e1003915.

Kovesi P (1999) Image features from phase congruency. Videre J Comput Vis Res 1:1–26.

Kriegeskorte N, Goebel R, Bandettini P (2006) Information-based functional brain mapping. Proc Natl Acad Sci U S A 103:3863–3868.

Kriegeskorte N, Kievit RA (2013) Representational geometry: integrating cognition, computation, and the brain. Trends Cogn Sci 17:401–412.

Kriegeskorte N, Mur M, Bandettini PA (2008) Representational similarity analysis-connecting the branches of systems neuroscience. Front Syst Neurosci 2:4.

Maris E, Oostenveld R (2007) Nonparametric statistical testing of EEG-and MEG-data. J Neurosci Methods 164:177–190.

Neider MB, Zelinsky GJ (2006) Scene context guides eye movements during visual search. Vision Res 46:614–621.

Nili H, Wingfield C, Walther A, Su L, Marslen-Wilson W, Kriegeskorte N (2014) A toolbox for representational similarity analysis. PLoS Comput Biol 10:e1003553.

Oliva A, Torralba A (2006) Building the gist of a scene: The role of global image features in recognition. Prog Brain Res 155:23–36.

Oostenveld R, Fries P, Maris E, Schoffelen J-M (2010) FieldTrip: open source software for advanced analysis of MEG, EEG, and invasive electrophysiological data. Comput Intell Neurosci 2011.

Peirce JW (2007) PsychoPy—psychophysics software in Python. J Neurosci Methods 162:8–13.

Recasens A, Khosla A, Vondrick C, Torralba A (2015) Where are they looking? In: Advances in Neural Information Processing Systems, pp 199–207.

Taulu S, Simola J (2006) Spatiotemporal signal space separation method for rejecting nearby interference in MEG measurements. Phys Med Biol 51:1759–1768.

Thompson KG, Hanes DP, Bichot NP, Schall JD (1996) Perceptual and motor processing stages identified in the activity of macaque frontal eye field neurons during visual search. J Neurophysiol 76:4040–4055.

Torralba A, Oliva A, Castelhano MS, Henderson JM (2006) Contextual guidance of eye movements and attention in real-world scenes: the role of global features in object search. Psychol Rev 113:766.

Van Essen DC, Dierker DL (2007) Surface-based and probabilistic atlases of primate cerebral cortex. Neuron 56:209–225.

Wandell BA (1995) Foundations of vision. Sinauer Associates.

Willenbockel V, Sadr J, Fiset D, Horne GO, Gosselin F, Tanaka JW (2010) Controlling low-level image properties: the SHINE toolbox. Behav Res Methods 42:671–684.

Yarbus AL (1967) Eye movements and vision. Springer.

